# Microphysiological engineering of the capillary interface of substantia nigra dopaminergic neurons to study vascular alterations in Parkinson’s Disease

**DOI:** 10.1101/2025.05.14.654131

**Authors:** Anika Alim, Yoongyeong Baek, Myungwoon Lee, Jungwook Paek

## Abstract

Parkinson’s Disease (PD) is primarily characterized by α-synuclein pathology, which manifests as intraneuronal inclusions, neuroinflammation, and neurodegeneration. However, emerging evidence also points to significant vascular impairments as a critical aspect of PD pathology, which remains largely underexplored due to the inability of traditional in vitro models to recapitulate such vascular changes. To address this unmet need, here we combine the human organ-on-a-chip technology with the principle of vasculogenic self-assembly to engineer the capillary interface of dopaminergic neurons in the substantia nigra pars compacta of the midbrain. In our proof-of-concept demonstration, we successfully recreated critical neuronal pathology in PD, including α-synuclein aggregation, inflammatory responses, and progressive neuronal degeneration, by exposing our model to specially generated PD-associated α-synuclein preformed fibrils. Importantly, this engineering approach also enables the investigation of progressive vascular changes characteristic of PD, such as endothelial dysfunction, barrier disruption, and vascular regression. Our sophisticated PD model establishes a novel platform for exploring the multifaceted nature of the disease and understanding the complex interplay between neurodegeneration and vascular pathology, offering a unique tool for developing innovative therapeutic strategies that address both the neuronal and vascular components of PD pathology.

## Introduction

Parkinson’s Disease (PD) is a debilitating neurodegenerative disorder that poses a significant clinical challenge worldwide^1–3^. It primarily manifests through hallmark motor impairments such as tremor, rigidity, and bradykinesia, which arise from the degeneration of dopaminergic neurons in the substantia nigra pars compacta (SNpc) of the midbrain^1,4–8^. Conventionally, research has focused on neuronal pathology within this specific brain area, exploring neuroinflammation, neurodegeneration, and eventual neuronal death^9–12^. However, recent insights have broadened our understanding, indicating that PD’s impact is not confined to the SNpc but extends to widespread neurodegenerative processes across multiple brain regions as the disease progresses^13–15^.

Central to this transmissive nature of PD is the accumulation of α-synuclein fibrils within affected neurons, which form neurodegenerative Lewy Bodies (LBs)^16–18^. These toxic aggregates exhibit prion-like behavior, templating the misfolding of native α-synuclein in healthy neurons and promoting the formation of new fibrils^14,19–24^. As the disease advances, this α-synuclein pathology spreads through interconnected brain regions, progressively amplifying neurodegeneration. Notably, even transplanted dopaminergic neurons are found to develop α-synuclein-rich LBs, indicating the potential for α-synuclein pathology to propagate across distinct brain areas^25,26^.

While neuronal proteinopathy is a defining feature of PD, there is growing recognition of associated vascular complications^27–34^. Clinical studies have revealed that PD patients frequently experience compromised vascular integrity and reduced cerebral perfusion, which may limit oxygen and nutrient delivery to neurons^35–38^. These vascular impairments may create a hostile neurovascular environment that exacerbates neuronal injury, suggesting that the vascular abnormalities could be key contributors to the progression of PD^29,39–41^. Despite its apparent pathophysiological significance, our understanding of the mechanisms underlying vascular dysfunction in PD remains incomplete. Recent animal studies have shown that misfolded α-synuclein including α-synuclein preformed fibrils (α-syn PFFs) is capable of inducing vascular abnormalities in the brain, establishing a link between α-synuclein pathology and vascular impairment^40,42–45^. Although these studies confirm vascular complications in PD and their association with α-synuclein aggregation, the inherent complexity of in vivo models complicates precise evaluations of how α-synuclein specifically impacts vascular structures and functions as the disease advances^46,47^. Consequently, these models often lack the resolution needed to dissect the dynamic interplay between vascular dysfunction and neurodegeneration in PD.

In an attempt to address the challenges of in vivo models, researchers have turned to in vitro models to study PD-associated vascular pathology under controlled conditions. Several studies using these models have demonstrated that α-syn PFFs may disrupt endothelial barrier integrity by compromising intercellular junctions in 2D endothelial monolayers, resulting in increased vascular permeability and compromised blood-brain barrier (BBB) function^28,44,48–51^. Therefore, this endothelial disruption implies that α-synuclein fibrils along with other neurotoxic agents, could more readily infiltrate brain tissues, potentially accelerating the spread and severity of α-synuclein pathology across diverse brain regions^52,53^. Although these reductionist models provide clear insights into the direct effects of α-synuclein fibrils on vascular endothelial cells, their typical use of 2D endothelial monolayers on porous polymeric membranes often fails to capture the complex 3D structure and physiological functions of the native neurovascular unit. As a result, the current in vitro models do not adequately represent the full scope of vascular pathology in PD, lacking the ability to accurately emulate crucial aspects such as structural changes in 3D capillary networks, disrupted blood perfusion, and the resulting compromised interactions between blood vessels and neurons. This gap in modeling capabilities poses a significant obstacle to advancing our understanding of the PD-associated vascular dysfunction and its mechanistic relationship with α-synuclein pathology.

Recognizing these technical limitations, we demonstrate how the synergistic integration of microengineering with biomedical strategies can be used to advance current PD modeling approaches. By leveraging human-organ-on-a-chip (HOOC) technology, we created a three-dimensional (3D) vascularized PD model that can precisely recapitulate the pathological effects of α-synuclein fibrils on both neurons and blood vessels, potentially allowing for an in-depth study of the progressive pathology at the neuronal-vascular interface in the SNpc (**Fig. 1a**). Specifically, we present a microfabricated platform that supports the self-organization of a vascular network through the vasculogenic process (**Fig. 1b**). By incorporating neuronal cells within this vascular bed, we constructed a 3D vascularized brain model that faithfully reproduces the specialized functions and dynamic structures of human midbrain tissue, including dopaminergic neurons with axonal and dendritic structures, as well as a perfusable vascular network. Following the creation of this model, we introduced α-synuclein fibril seeds which were generated under the disease-associated membrane conditions into the microengineered device to mimic PD in vitro. This approach successfully recapitulates key pathological features of PD, including the intraneuronal formation of α-synuclein-rich fibrils, neuroinflammation and neurodegeneration. As a proof-of-concept demonstration, we utilized this disease modeling approach to investigate the impacts of α-synuclein fibrils on the structure and function of the 3D vascular network. Our multidisciplinary engineering strategy thus establishes a pathophysiologically relevant platform, opening new avenues to investigate the multifaceted nature of PD and to deepen our understanding of the complex interplay between neuronal α-synuclein pathology and vascular dysfunction.

**Figure 1.**
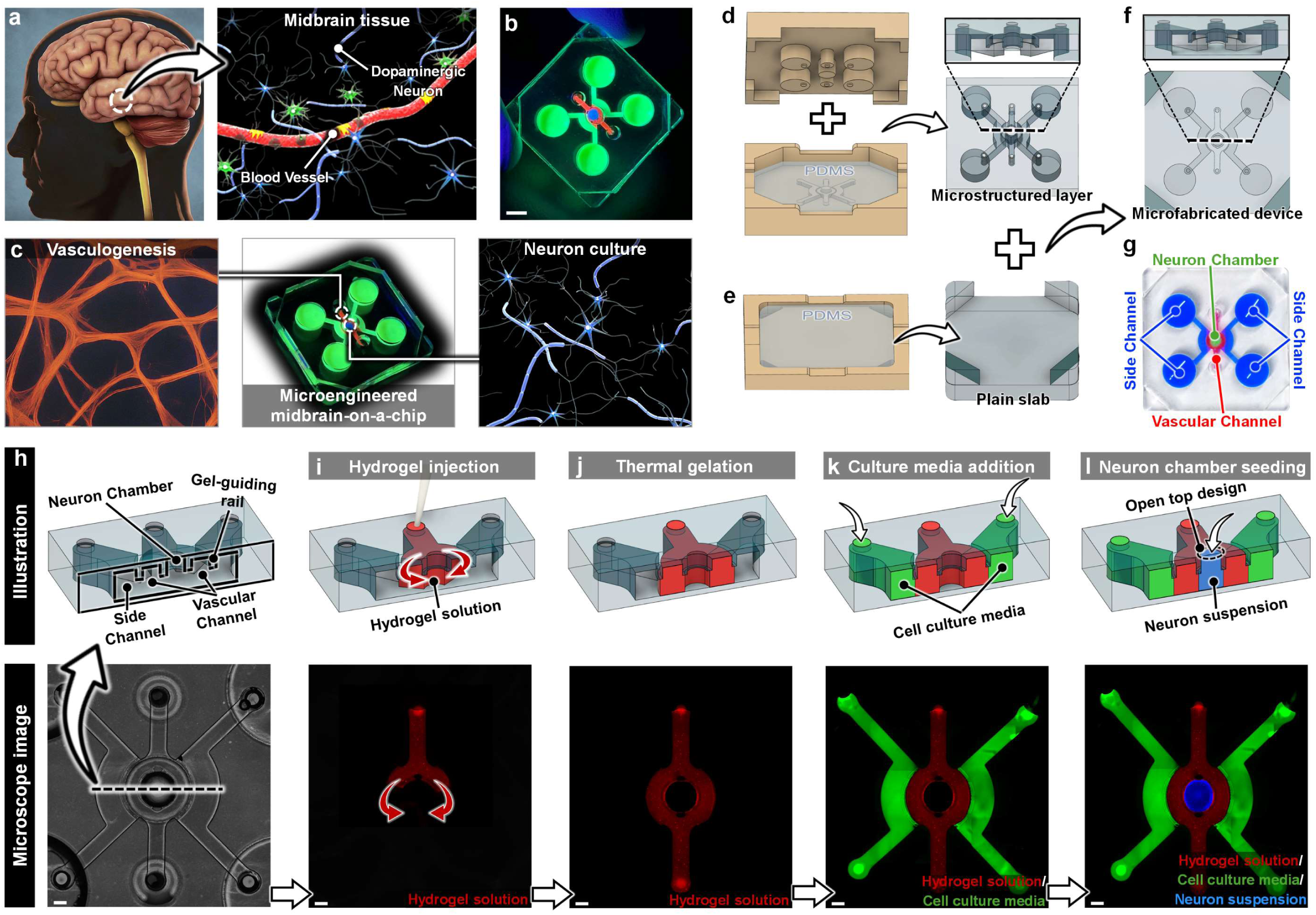
Microengineered 3D culture platform to emulate the capillary-tissue interface of the human midbrain. **a**, Human midbrain tissue featuring dopaminergic neurons and a vascular network. **b**, Microfabricated culture platform designed to engineer vascularized midbrain tissue in vitro. Scale bar, 1 mm (photo). **c,** Schematic illustration of the microengineered midbrain-on-a-chip integrating a vascular network with neuron culture. **d-f,** Fabrication process using 3D printed molds to create a microstructured PDMS replication (**d**) and a plain PDMS slab (**e**). The assembly of these two PDMS layers forms the midbrain tissue culture platform (**f**). **g,** Assembled device features the neuron chamber (green), the vascular channel (pink), and the two side channels connected to reservoirs (blue). **h,** Cross-sectional view of the assembled device shows that the neuron chamber, vascular channel and side channels are partially separated by gel-guide rails. **i-j,** Sequential process for constructing multiple tissue compartments begins with the injection of a hydrogel solution (red) into the vascular channel, shown without spillage in the fluorescent microimage (bottom) (**i**). Thermal gelation then solidifies the hydrogel compartment (red) within the vascular channel, as demonstrated in the bottom fluorescent microimage (**j**). To support cell culture within the vascular channel’s hydrogel scaffold, a media solution (green) is subsequently added to the side channels (**k**). Additionally, a solution containing neuronal cells is introduced into the neuron chamber through its open-top to establish in vitro neuronal tissue (**l**). Scale bars, 50 µm.

## Result and Discussion

### Platform Design and Construction

Considering the anatomical features of midbrain tissues affected by PD, our microengineered culture platform was designed to enable in vitro reproduction of physiological tissue compartmentalization of the midbrain, which includes dopaminergic neurons and vasculature while maintaining their essential interactions for a comprehensive study of PD progression. To achieve this, our system contains a central neuron chamber for culturing midbrain dopaminergic neurons (**Fig. 1c**). Surrounding this compartment, additional microchannels are dedicated to the vasculogenic production of 3D microvascular networks and providing the necessary media support for multiple tissue cultures within our device (**Fig. 1c**).

This microdevice was fabricated using standard soft-lithography techniques. The fabrication process began with two specially designed 3D molds, one featuring microstructures for a central neuron chamber and surrounding microchannels, and the other for cell culture media reservoirs, as illustrated in **Fig. 1d**. Between these molds, liquid phase PDMS was introduced and, once cured, it replicated the microstructures of the 3D molds. As a result, the PDMS replication comprises an open-top neuron chamber and surrounding microchannels on one side, and cell culture media reservoirs on the opposite side (**Fig. 1d**). This microstructure-featured PDMS layer was then assembled with a plain PDMS layer, which was soft-lithographically generated using its corresponding mold (**Fig. 1e**), resulting in the creation of the microengineered culture platform depicted in **Fig. 1f**. The different color codes in **Fig. 1g** distinctly indicate the neuron chamber (green), the surrounding vascular channel (red), and the feeding side channels (blue), confirming their respective locations within our microengineered device.

Within our microengineered device, the neuron chamber, vascular channel, and two side channels are designed to achieve compartmentalization ultimately (**Fig. 1h**), but initially, they are only partially separated by thin PDMS rail-structures. These thin rail structures extend from the top surface of the device downward but do not reach the bottom (**Fig. 1h**), thus allowing connectivity between the compartments at their base. Strategically positioned at the interfaces between the vascular channel and the side channels, as well as between the vascular channel and the neuron chamber, these PDMS rails play a crucial role in constructing a hydrogel scaffold that contains a 3D vascular network within the vascular channel. This approach enables the eventual formation of distinct, yet functionally integrated neuron chambers and side channels on both sides of the vascular channel. Specifically, when a cell-embedding hydrogel solution is injected into the vascular channel, the PDMS rails guide the hydrogel’s movement by leveraging the surface tension between the hydrogel, the rails, and the device’s bottom surface to confine it within the vascular channel. This process is demonstrated by the directed flow of the hydrogel solution (red) along the vascular channel in our device which remains confined without spilling over into the neuron chamber and side channels as shown in **Fig. 1i**. Following the gel injection, subsequent thermal gelation solidifies this arrangement, creating a cell-laden hydrogel scaffold that defines the structural boundaries between the neuron chamber and side channels (**Fig. 1j**). Consequently, this hydrogel scaffold effectively establishes distinct and functional compartments for the neuron chamber and the flanking side channels. Cells are then cultured within the vascular channel hydrogel with media supplied through the side channels (**Fig. 1k**). Notably, the neuron chamber features an open-top design, enabling the modular incorporation of various types of cells into this hollow chamber. Thus, neuronal cells are seeded into the neuron chamber through its open top and cultured to model vascularized neuronal tissues (**Fig. 1l)**. The completion of this process yields a 3D tissue construct where neuronal tissue functionally interacts with the vasculature.

### Construction of engineered human midbrain tissue

In our first study, we aimed to demonstrate the capability of our tissue production approach described above for reconstructing the capillary interface within the SNpc, a midbrain region critically impacted in PD. To this end, we started with the construction of a vascularized hydrogel scaffold within the vascular channel of our microengineered device as depicted in **Fig. 2a**. This was achieved by embedding both human umbilical vein endothelial cells (HUVECs) and normal huma lung fibroblasts (NHLFs) into a fibrinogen solution, which was simultaneously mixed with thrombin to transform the hydrogel solution into the cell-laden fibrin gel scaffold. Before gelation, thus we then quickly injected the cell-containing fibrinogen solution into the vascular channel (**Fig. 2b**), where gelation occurred, forming a stable fibrin gel scaffold.

**Figure 2.**
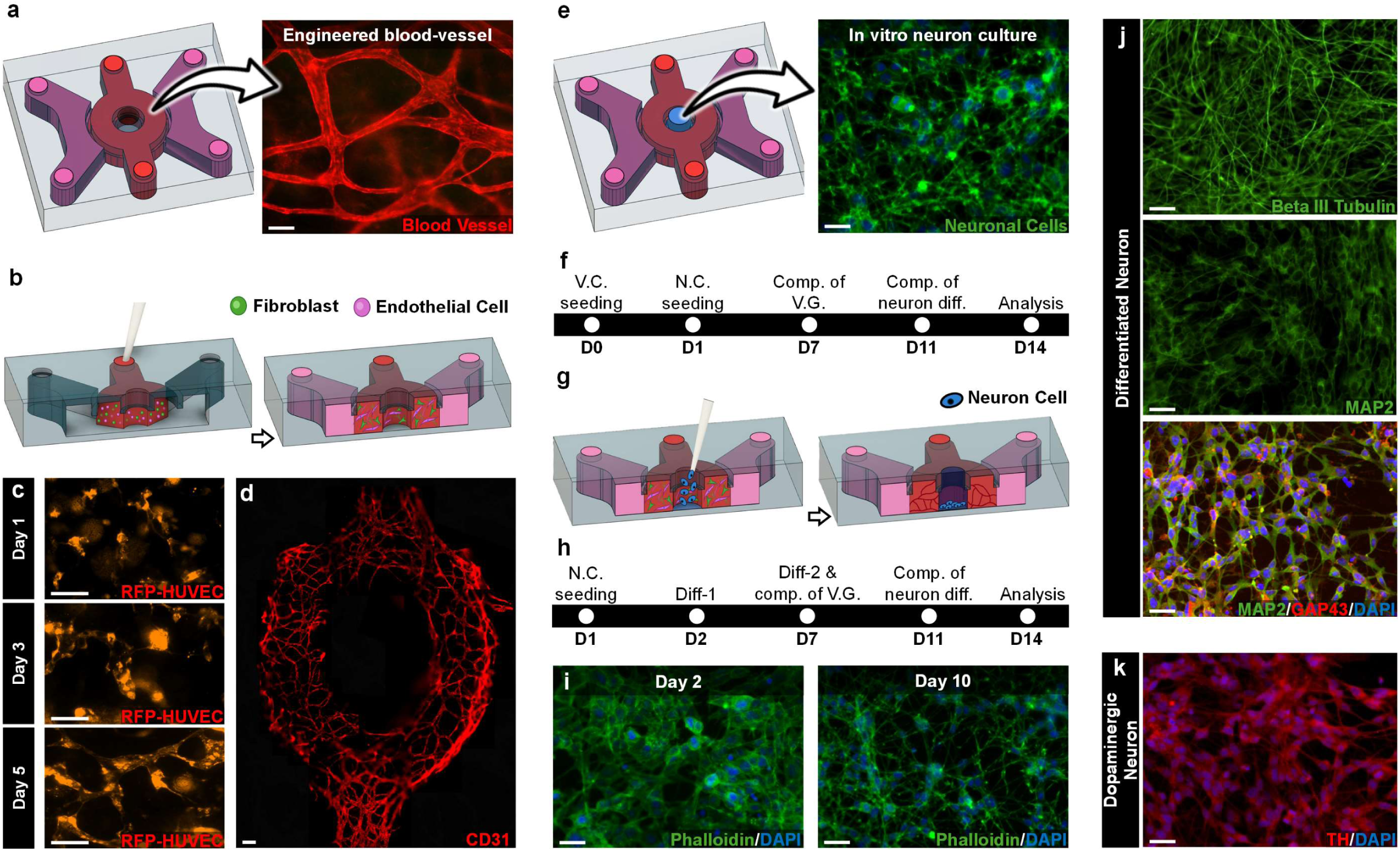
Co-culturing of 3D vascular network with dopaminergic neuronal cells in the human midbrain-on-a-chip. **a,** Schematic illustration of the microengineered device containing a vascular network within its vascular channel. **b,** To form vasculature, endothelial cells and fibroblasts suspended in an extracellular matrix (ECM) hydrogel solution are injected into the vascular channel (left). Following gelation, endothelial cell growth media is added into the side channels to support vasculogenesis (right). **c,** Over time, the coculture of RFP-HUVECs (red) and fibroblasts (not shown) in the hydrogel results in the formation of network structures. Scale bars, 200 µm. **d,** Immunofluorescence staining demonstrates CD31 expression by the endothelial tubes in the self-assembled vascular network within the vascular channel. Scale bar, 50 µm. **e,** Schematic representation of neuronal tissue reproduction in the neuron chamber. **f,** Timeline of the coculture of neuronal cells and vasculature in the device. Abbreviations: V.C. (vascular channel), N.C. (neuron chamber), V.G. (vasculogenesis), D (day), comp.(completion), and diff. (differentiation). **g,** Schematic diagram showing neuroblastoma cell seeding into the neuron chamber through its open-top design. **h,** Timeline of dopaminergic differentiation in neuroblastoma cells. **i,** Phalloidin staining reveals morphological changes in neuroblastoma cells during differentiation on Day 2 and Day 10. Scale bars, 100 µm. **j,k,** Micrographs show the successful differentiation of neuroblastoma cells into dopaminergic neuronal cells after 11 days of co-culture. β III tubulin (green, top), MAP2 (green, middle) and GAP43 antibodies stain the neuronal cytoskeleton, dendritic structures, and axonal projections, respectively (**j**). Immunostaining for tyrosine hydroxylase (TH, red) confirms dopaminergic neuron differentiation (**k**). Scale bars, 100 µm.

Following the thermal gelation, the cells were uniformly dispersed throughout the gel and began to spread within 24 hours of seeding, as evidenced by the use of red fluorescent protein (RFP)-expressing HUVECs (Day 1 in **Fig. 2c**). During culture, microfluorimetric imaging of the embedded RFP-HUVECs revealed the development of network structures within the fibrin gel. As early as on day 3, HUVECs appeared significantly elongated and began to organize into chord-like structures (Day 3 in **Fig. 2c**). These morphological changes became more pronounced over time and eventually led to the formation of cellular networks that resembled the primitive vascular plexus generated by vasculogenesis in vivo (Day 5 in **Fig. 2c**). Further validating our vasculogenic approach, immunostaining analysis at Day 7 confirmed the formation of intricate networks of interconnected endothelial cells that exhibited robust expression of cluster of differentiation 31 (CD31) throughout the vascular channel as shown in **Fig. 2d**. Notably, this expression was not observed in the central region of our device, which is dedicated to neuronal tissue production (**Fig. 2d**). Therefore, the spatially specific configuration of CD31 expression of the engineered vascular 45 network demonstrates the efficacy of our tissue culture platform in recapitulating biological vascularization processes and ensuring the compartmentalization of the vascular channel and neuronal chamber which is crucial for the simultaneous yet separate development of vascular and neuronal tissues within our microengineered device.

In the next phase of our study, we leveraged the capabilities of our microengineered tissue culture platform to develop in vitro models of vascularized midbrain tissues, essential for PD modeling. As a first step towards this end, neuroblastoma cells were introduced into the neuron chamber of our device with the aim to differentiate them into dopaminergic neurons characteristic of the SNpc, which are primarily affected in PD (**Fig. 2e**). This generation of midbrain neurons was conducted concurrently with the vasculogenic production of engineered vessels within the vascular channel (**Fig. 2f**), ensuring the formation of an integrated network that closely interacts with the neuronal tissue component. Specifically, one day after the onset of the vasculogenic culture, neuroblastoma cells were introduced into the neuron chamber and maintained until they reached approximately 100% confluency, in parallel with vasculature formation within the vascular channel (**Fig. 2f**). It should be noted that the open-top design of the neuron chamber not only facilitates the modular integration of neuronal tissues directly into the engineered vascular beds to mimic the interface between blood vessels and brain parenchymal tissues in a tissue-specific manner but also enables the direct supply of medium to the cultured neuronal cells (**Fig. 2g**). Once the neuroblastoma cells reached confluence, their growth media was switched to two types of differentiation media, each specifically formulated and applied sequentially to induce their transformation into dopaminergic neuron types (**Fig. 2h**). This differentiation protocol is based on previous studies demonstrating that neuroblastoma cells are capable of developing synaptic connections and synthesizing dopamine as a neurotransmitter^54^. Consistent with these findings, the neuroblastoma cells in our device began to extend multiple thin cellular projections during the early stages of differentiation (Day 2 in **Fig. 2i**) and developed complex axonal and dendritic structures over time, indicative of their potential to form synaptic connections (Day 10 in **Fig. 2i**).

These cellular projections were positively stained for β III tubulin, MAP2, and GAP43, markers specific for the neuronal cytoskeleton, dendritic structures, and axons, respectively (**Fig. 2j**), confirming the successful differentiation of neuroblastoma cells into neuronal cells. The dopaminergic capability of the differentiated neuronal cells was also demonstrated through immunofluorescent staining for tyrosine hydroxylase (TH), essential enzyme for dopamine biosynthesis and predominantly found in dopaminergic neurons of the midbrain’s SNpc (**Fig. 2k**). Notably, as dopaminergic neurons were generated, HUVECs and NHLFs embedded within the fibrin gel scaffold of the vascular channel formed a vascular construct without compromising the viability and differentiation of the dopaminergic neurons in the neuron chamber (data not shown).

While this study demonstrates the feasibility and potential of our engineering approach to model vascularized midbrain tissues, it is important to note that there remains significant room for further studies to improve the fidelity of our microengineered midbrain tissues. Cellular heterogeneity and specificity, critical features at the capillary interface of the SNpc have not yet been fully recapitulated in our model. In particular, it is possible to incorporate brain pericytes that are intimately associated with the endothelial lining of the microvasculature within brain tissues and serve as a key regulator of vascular structure and function in a variety of physiological processes ^55,56^. Similarly, introducing astrocytes that are prevalent in the perivascular region and play a key role in supporting physiological functions of dopaminergic neurons and cerebrovasculature, may also enhance the physiological relevance of our model^57,58^. Thus, further research efforts may be necessary to engineer the intrinsic cellular characteristics of the midbrain with the goal of recapitulating a more physiologically relevant cellular microenvironment, which may be crucial for studying the effects of a multi-cellular environment on the spread of diverse brain disease conditions. Nevertheless, our study indicates the capabilities of our microengineered tissue culture platform in emulating the physiologically relevant structures and functions of dopaminergic neurons interfaced with a vascular network, which has been challenging to achieve with conventional 2D cell culture models.

### Generation of synucleinopathic neuronal conditions in the vascularized midbrain model

α-synuclein is a major component of intraneuronal fibrillar inclusions known as Lewy bodies (LBs), a hallmark of PD and other related synucleinopathies^16–18^. As the disease progresses, native α-synuclein undergoes pathological misfolding and aggregation, triggering neuronal oxidative stress, neuroinflammation, and synaptic dysfunction which are key pathological processes that drive neurodegeneration, ultimately leading to neuronal loss^59–66^. Therefore, the ability of our vascularized midbrain model to accurately capture this α-synuclein pathology is essential for its application as a PD modeling platform, particularly for investigating disease progression and its impact across distinct brain tissue constructs. Recognizing this need, we next explored whether our midbrain model could be leveraged to develop a specialized in vitro brain disease model capable of recapitulating the α-synuclein pathology in PD.

For this investigation, we utilized short segments of α-synuclein fibrils known as α-syn PFFs as a pathological seed, chosen for their well-established neurotoxicity and ability to drive α-synuclein pathology in PD^21,67–70^. Previous studies typically employed α-syn PFFs prepared without cellular components in buffers to explore their pathological effects in PD^67,71^. However, growing evidence suggests that interactions between α-synuclein and cellular lipid membranes during aggregation critically modulate the conformational shape of α-synuclein fibrils^72–75^, which may determine their development and propagation as PD progresses^76–78^. Motivated by this insight, we first generated specialized α-syn PFFs by emulating the lipid composition of neuronal plasma membranes affected by PD^72,79,80^, which consist of sphingomyelin, DPPC (1,2-dipalmitoyl-sn-glycero-3-phosphocholine), POPE (1-palmitoyl-2-oleoyl-sn-glycero-3-phosphoethanolamine), and cholesterol during α-synuclein fibril formation as depicted in **Fig. 3a**. Once these PD-associated α-syn PFFs were generated, we then exposed our vascularized midbrain model to the fibril seeds through the open-top of the neuron chamber with their concentration of 1.0 μM to induce key pathological features observed in dopaminergic neurons in PD as illustrated in **Fig. 3b**.

**Figure 3.**
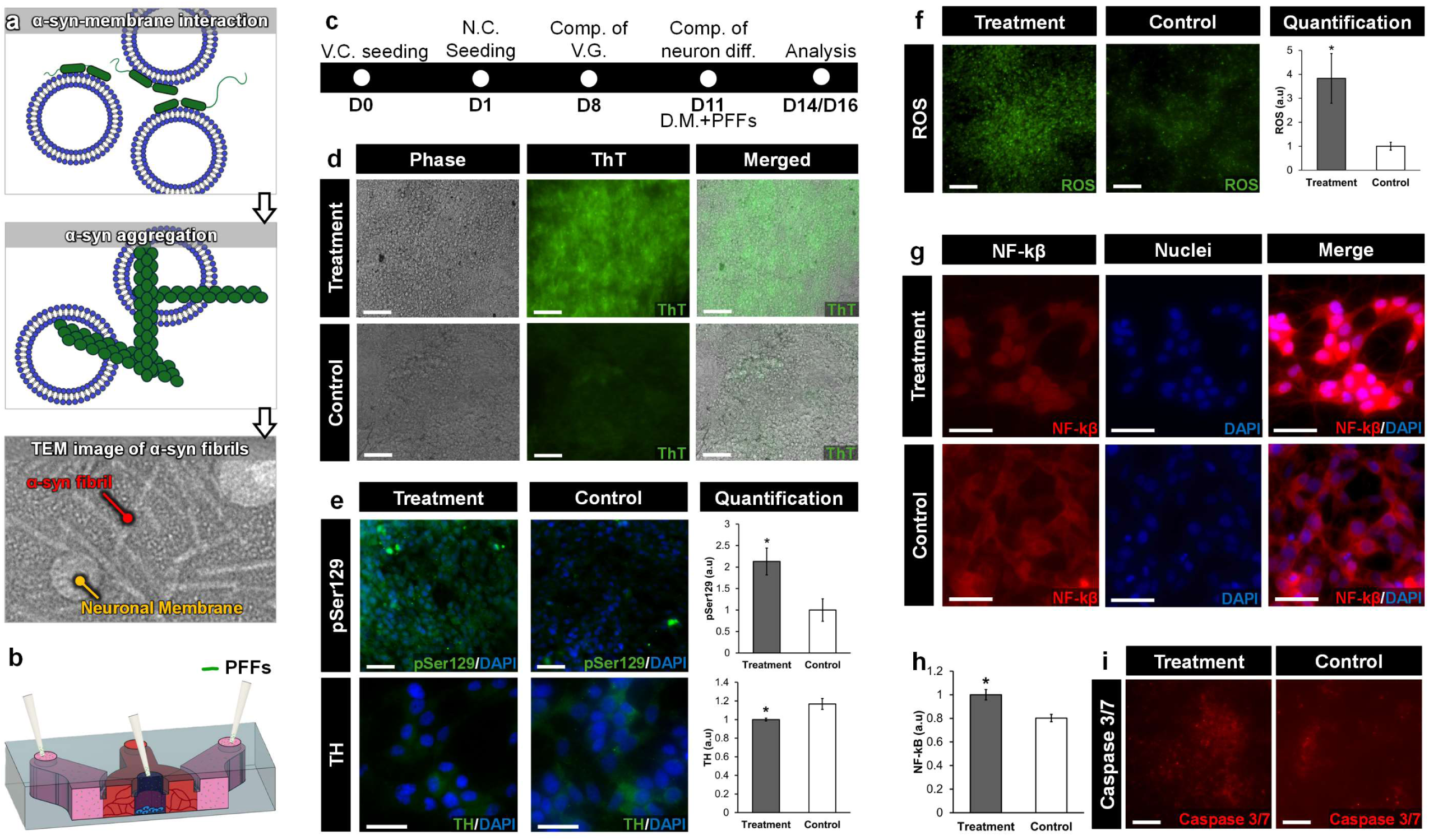
Microphysiological modeling of α-synuclein-induced neurodegeneration in engineered midbrain tissue. **a,** Sequential process for generating pathologically relevant α-synuclein preformed fibrils (PD-associated α-syn PFFs) using lipid vesicles that mimic the lipid composition of PD-affected neurons. TEM (Transmission Electron Microscopy; bottom) image shows the successful production of short form α-synuclein fibrils. **b,** Treatment of dopaminergic neuronal cells with PD-associated α-syn PFFs to emulate neurotoxicity in our midbrain model. **c,** Timeline of PD-associated α-syn PFF treatment and analysis. ‘D.M’ denotes the differentiation media used for neuronal cells. **d,** Thioflavin T (ThT) fluorescence confirms the formation of α-synuclein fibrils templated by internalized PD-associated α-syn PFFs in dopaminergic neuronal cells. Scale bars, 100 µm. **e,** Immunostaining analysis of phosphorylated α-syn (pSer129) and TH indicates increased misfolding of endogenous α-synuclein and reduced TH expression following PD-associated α-syn PFF treatment. Scale bars, 100 µm and 200 µm, respectively. **f,** Production of reactive oxygen species (ROS) is significantly elevated in response to PD-associated α-syn PFF treatment. Scale bars, 100 µm. **g, h,** Dopaminergic neuronal cells subjected to PD-associated α-syn PFFs also exhibit activation and nuclear translocation of NF-κB (**g**). Quantification of NF-κB translocation further demonstrates the activation of the transcriptional factor (**h**). Scale bars, 200 µm. **i,** Exposure to PD-associated α-syn PFFs induces neuronal apoptosis as evidenced by the increased expression of Caspase-3/7 compared with control. Scale bars, 100 µm. ***P < 0.001, **P < 0.01, *P < 0.05. Data show mean ± SD with n = 3.

Treating our device with the PD-associated α-syn PFFs for 2 days (**Fig. 3c**) significantly promoted fibrilization within the dopaminergic neuronal cells as demonstrated in **Fig. 3d**. Compared to the untreated control group, this α-syn PFF treatment resulted in a two-fold increase in Thioflavin T (ThT) fluorescence intensity, a dye commonly used to probe intracellular protein fibrils, indicating that our PD-associated α-syn PFFs effectively induce intraneuronal fibril aggregation, a defining pathological feature of PD (**Fig. 3d**). After the administration of PD-associated α-syn PFFs, we allowed our devices a 3-day stabilization period (**Fig. 3c**). This step was crucial to let the fibrils fully interact with the cellular environment, establishing the experimental conditions necessary for a robust evaluation of their pathological impact. We then assessed how these fibrils influenced α-synuclein pathology within our disease model. For this investigation, our analysis initially focused on the PD model’s ability to recapitulate the phosphorylation of native α-synuclein, indicative of the protein misfolding which is a critical pathological process in the prion-like spread of α-synuclein pathology as PD progresses^81–84^. Compared to the control group, the dopaminergic neuronal cells within devices treated with PD-associated α-syn PFFs displayed more than 2 times higher levels of α-synuclein phosphorylated at the serine-129 site, a dominant yet abnormal modification found in Lewy bodies (LBs)^81–84^ as shown in **Fig. 3e**. Our immunofluorescence analysis also revealed significant alterations in TH expression in these treated neuronal cells as evidenced in **Fig. 3e**. Consistent with previous in vivo studies^84–89^, the α-synuclein PFF treatment led to a remarkable decrease in TH expression within our midbrain models, measured as a 20% decrease (**Fig. 3e**). This decrease suggests abnormalities in dopamine synthesis potentially linked to PD, which may disrupt dopamine homeostasis in the SNpc and thus impair its critical role in motor control.

Growing evidence indicates the critical role of neuroinflammation in the progression of neurodegenerative diseases including PD^59–66^. Building upon this insight, the subsequent phase of our study evaluated the capability of our midbrain model to recapitulate neuronal responses linked to neuroinflammation. Specifically, we first investigated the production of reactive oxygen species (ROS) by the dopaminergic neuronal cells in response to the PD-associated α-syn PFFs. This aspect of the study was chosen for examination as a proinflammatory factor because α-synuclein fibrils have been demonstrated to significantly enhance neuronal ROS production by activating reduced nicotinamide adenine dinucleotide phosphate (NADPH) oxidases, thereby driving oxidative stress within neurons^59,90,91^. Therefore, our experiments focused on measuring neuronal ROS production in response to the PD-associated α-syn PFFs, aiming to validate our model’s capacity to emulate this key pathological process in PD. Consistent with previous results^66,90^, the treatment with the PD-associated α-syn PFFs in our model resulted in a 4-fold increase in ROS levels compared to the control group as displayed in **Fig. 3f**. Given the established role of ROS as key mediators of neuroinflammation^62,93,94^, we further examined whether the PD-associated α-syn PFFs also elicited inflammatory responses in the dopaminergic neuronal cells. For this study, we focused on the analysis of nuclear factor-kappa B (NF-κB) as a representative readout of neuroinflammation considering it is a transcription factor known to regulate genes involved in inflammation and stress responses^95–97^. In the control devices shown in **Fig. 3g**, immunostaining showed that NF-κB was primarily localized in the cytoplasm of dopaminergic neuronal cells, indicating its inactive state. However, following treatment with PD-associated α-syn PFFs, a noticeable shift in NF-κB distribution occurred with a significant increase in nuclear immunofluorescence intensity and a corresponding decrease in cytoplasmic levels (**Fig. 3g**). This change was indicative of the activation and nuclear translocation of NF-κB triggered by the treatment of PD-associated α-syn PFFs. Quantitative analysis of this observation revealed that exposure of the dopaminergic neuronal cells to the PD-associated α-syn PFFs resulted in a 2-fold increase in nuclear translocation of NF-κB compared to the control (**Fig. 3h**). While further research is necessary, these findings suggest that the pathological responses of our engineered dopaminergic neuronal tissues to PD-associated α-syn PFF treatment are involved in the development of neuroinflammation.

It was noted that treatment with PD-associated α-syn PFFs resulted in a reduction of dopaminergic neuronal cells within our devices (**Figs. 3e,g**), indicating the neurotoxic impact of the pathological agents. Given that neuroinflammation can trigger a cascade of processes leading to neuronal apoptosis^97–99^, we extended this observation to determine whether the PD-associated α-syn PFFs specifically activate apoptotic pathways, a known hallmark of dopaminergic neuronal loss in PD pathology. Therefore, our subsequent experiments were focused on measuring the activation of caspase-dependent apoptotic pathways by quantifying caspase-3/7 expression in the neuronal cells treated with PD-associated α-syn PFFs. As shown in **Fig. 3i**, exposure to PD-associated α-syn PFFs led to a pronounced increase in caspase-3/7 expression compared to the control group, suggesting a robust activation of apoptosis in the treated neuronal cultures. These data suggest that α-synuclein fibrils may trigger programmed cell death^100,101^, which could explain the dopaminergic neuronal loss observed in α-syn PFF-treated devices and potentially offer insights into similar neurodegenerative processes in the brains of PD patients.

Taken together, our studies demonstrate the capabilities of our PD modeling approach in emulating the cascade of processes implicated in α-synuclein pathology within dopaminergic neurons, including neurodegeneration, neuronal oxidative stress, neuroinflammation, and progressive neuronal death. It remains to be examined, however, how the responses of other brain cell populations, such as microglial cells and astrocytes, interact with those of dopaminergic neurons in the presence of α-synuclein fibrils, which is an important consideration for more accurately recapitulating and examining the development and propagation of α-synuclein pathology. Nevertheless, the data presented here demonstrates the potential of this neurodegenerative disease model to simulate and investigate the intraneuronal manifestations of PD.

### Microphysiological model of α-synuclein-involved vascular impairment

Recent studies increasingly point to the crucial role of vascular pathology in PD, identifying various vascular alterations that emerge and possibly aggravate neurodegenerative processes^27–34^. These vascular changes include increased BBB leakage, vascular regression, and disruptions in cerebral blood flow, which collectively contribute to neuronal injury and accelerate the disease progression^27,31,33,34,44,103–106^. Particularly at the capillary interface of dopaminergic neurons in the SNpc, such vascular dysfunction may compromise nutrient and oxygen delivery, hinder effective drug transport, and promote the accumulation of neurotoxic substances in the immediate vicinity of neurons^31,34,38,41^. Additionally, these vascular impairments may alter cellular signaling within the neurovascular unit, further disrupting the neuronal microenvironment and potentially intensifying the progression of PD^29,39^. Notably, evidence from animal models suggests that these vascular changes can occur before significant neuronal loss, implicating them in the early pathogenesis of PD^31,106,107^. Despite their significant pathophysiological implications in PD, the precise mechanisms underlying these vascular changes remain elusive. Emerging evidence has identified α-synuclein fibrils as key factors in increasing BBB permeability^28,44,48–51^, yet a comprehensive understanding of their role in the broader spectrum of PD-associated vascular pathology is still lacking. This knowledge gap persists largely due to the limited capacity of existing PD models to fully replicate the complex and progressive neurovascular abnormalities observed in the neurodegenerative disease.

To suggest an alternative in vitro strategy to tackle this technical deficiency, we explored whether our vascularized midbrain model could be leveraged to emulate the complex vascular changes observed in PD. Recognizing the well-established role of α-synuclein fibrils in compromising BBB integrity^28,44,48–51^, the initial phase of our study focused on applying PD-associated α-syn PFFs to assess their effects on both the structural and functional integrity of the vasculature within our microengineered device. To this end, we introduced the PD-associated α-syn PFFs into two side channels interfacing with the engineered vascular network which was established through the vasculogenic interactions of endothelial cells with fibroblasts (**Fig. 4a**). For this work, we employed a 5.0 µM concentration of PD-associated α-syn PFFs, which was five times higher than that used for dopaminergic neuronal cells discussed in the previous section. This higher concentration was chosen because the 3D fibrin gel scaffold might impede the diffusion of the fibrils, potentially mitigating their impact on the embedded vasculature. After 2-day treatment with the α-syn PFFs, the conditioned media was replaced with fresh media for both treated and control groups which were then allowed to rest for an additional 3 days before carrying out analyses, as detailed in **Fig. 4b**.

**Figure 4.**
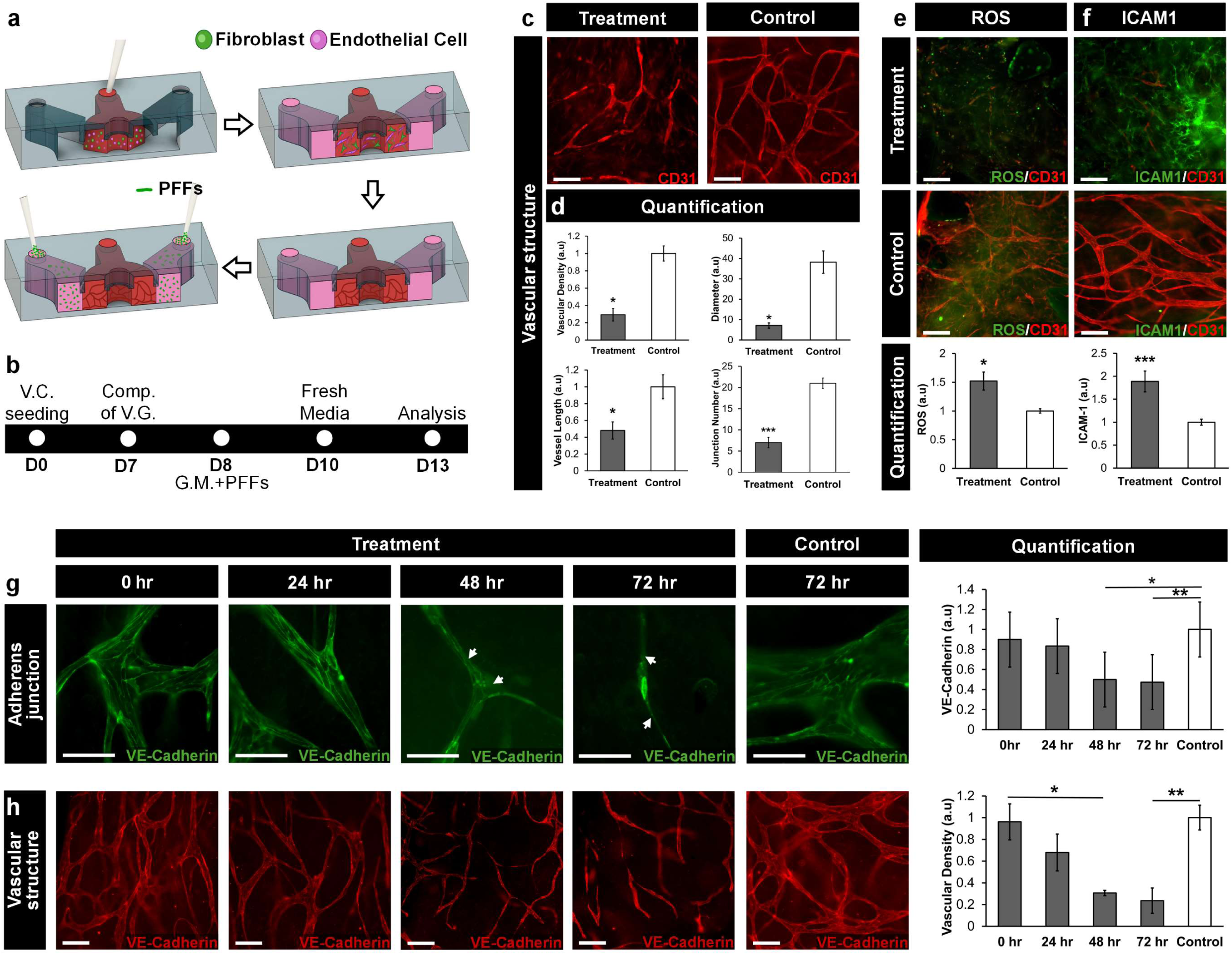
Microengineering of pathological vascular alterations in PD. **a,** Schematic illustration of exposing the 3D vascular network in our device to PD-associated α-syn PFFs. b, Timeline of PD-associated α-syn PFF treatment and analysis. ‘G.M.’ denotes endothelial growth media. **c,d,** Our vascularized midbrain model allows for the detection and analysis of vascular toxicities induced by PD-associated α-syn PFFs. The fibrils cause the loss and disintegration of vasculature (**c**), resulting in significant reductions in vascular density, average vessel diameter, total vessel length, and junction number (**d**). Scale bars, 100 µm. **e,** Exposure to PD-associated α-syn PFFs induces oxidative stress of engineered vessels within our device. Scale bars, 100 µm. **f,** The deleterious potential of PD-associated α-syn PFFs to elicit vascular inflammation is demonstrated by increased endothelial expression of ICAM-1. Scale bars, 100 µm. **g,** Immunofluorescence analysis of VE-cadherin at various time points (0, 24, 48, and 72 hours) shows progressive disruption of endothelial cell junctions due to treatment with PD-associated α-syn PFFs. Scale bars, 100 µm. **h,** VE-cadherin staining also reveals detrimental changes in vascular structures over time following treatment with PD-associated α-syn PFFs. Scale bars, 100 µm. ***P < 0.001, **P < 0.01, *P < 0.05. Data show mean ± SD with n = 3.

With the vasculature thus treated, we proceeded to assess the vascular responses using microfluorimetric techniques to examine changes across a range of phenotypic endpoints indicative of both vascular structure and biochemical function. Considering the characteristic vascular regression observed in the SNpc of PD patients^27,33,34,38^, our initial analysis focused on examining morphological alterations within the vasculature of treated and control groups. As shown in **Fig. 4c**, the exposure to the PD-associated α-syn PFFs led to pronounced structural changes, evidenced by the significant disruption of its characteristic network architecture. The detrimental effect of the α-syn fibrils was further evidenced by the regression of the interconnected tubular network to disorganized endothelial chord-like structures as well as a notable decrease in vascular density (**Fig. 4c**). Our quantitative analysis confirmed these observations, revealing a significant reduction in vascular density due to extensive loss of endothelial lining in vessels treated with the PD-associated α-syn PFFs (**Fig. 4d)**. Additionally, further estimates of vascular architecture showed significant decreases in average vessel diameter, total vessel length, and the number of vessel junctions over the same period (**Fig. 4d**). These results are consistent with previously observed vascular regressions in PD patients, which are characterized by endothelial degeneration, reduced vessel length and branching points, and increased vessel diameter^27,33^.

Despite these clinical observations, little progress has been made in emulating and investigating the complex vascular changes associated with PD, thereby limiting our understanding of the fundamental processes underlying the progression of this pathological feature. Thus, our findings indicate the potential of our engineering approach to bridge the gap in understanding PD-associated vascular pathology, particularly by delineating how blood vessels are affected by α-synuclein fibrils in PD.

Following our investigation into the structural impact of PD-associated α-syn PFFs on vascular pathology, we next sought to examine molecular mechanisms potentially implicated in the vascular regression. Specifically, we analyzed the production of ROS and the expression of intercellular adhesion molecule-1 (ICAM-1), both of which play key roles in inflammatory and immune-mediated endothelial dysfunction, potentially leading to adverse vascular responses^93,109,110^. Consequently, our immunofluorescence analysis revealed that treatment with PD-associated α-syn PFFs led to a 1.5-fold increase in ROS levels compared to the control group (**Fig. 4e**). This observation was accompanied by a cascade of inflammatory responses, as evidenced by an approximately 2-fold increase in ICAM-1 expression (**Fig. 4f**).

Building upon these observations, we further explored the downstream effects of the PD-associated α-syn PFFs on vascular barrier integrity. Specifically, our focus turned to the adherens junction protein, Vascular Endothelial Cadherin (VE-cadherin) which serves as a crucial marker of vascular integrity^110^. Given the emerging evidence of a dynamic progression in PD associated vascular changes which may start from subtle BBB leakage to eventual vascular regression^31^, our study also explored whether our engineering approach could emulate these temporal dynamics. To this end, we analyzed the expression of this adherens junction protein at specific intervals (0, 24, and 48 hours) during our standard 2-day treatment period with PD-associated α-syn PFFs. Additionally, to capture more comprehensive data on the progression of vascular changes, we extended our observations to 72 hours, allowing for an extended assessment period, and then compared the outcomes with those in the control group. As shown in **Fig. 4g**, our immunostaining results revealed adherens junction damage in all devices treated with the PD-associated α-syn PFFs. This damage progressively worsened over time, starting with a slight decrease in VE-cadherin expression at 24 hours. By 72 hours, the junctional disruption became more severe, as evidenced by a significant loss of VE-cadherin expression and the absence of clearly defined staining at endothelial cell junctions (indicated by arrowheads in **Fig. 4g**). Consistent with these observations, the α-syn PFF treatment progressively altered vascular structures, evident from mild vessel thinning at 24 hours with negligible structural disruption, and eventually resulting in significant vessel disintegration and decreased vascular density by 72 hours (**Fig. 4h**). These data demonstrate the temporal dynamics of vascular changes under the influence of α-syn PFF treatment, which may begin with subtle endothelial barrier disruptions and intensify to complete vascular disintegration as PD progresses. It should be noted that following 48 hours of treatment, our device exhibited a substantial increase in the number of thin and acellular vessels, reminiscent of string vessels typically seen in PD patients^33,34^. This observation suggests that our approach provides a valuable platform for investigating the neurodegenerative impact of these non-functional capillary remnants that remain challenging to model in vitro. In contrast, untreated control devices exhibited negligible changes in VE-cadherin expression and vascular structure during the same period.

Collectively, our findings suggest that α-synuclein fibrils exert cytotoxic effects on vascular endothelial cells, potentially inducing oxidative stress, endothelial disruption, and chronic inflammation, which may collectively contribute to vascular injury. Beyond their role in neurodegeneration, these results indicate that α-synuclein fibrils play a significant role in PD-associated vascular pathology, providing insights into how they may contribute to early endothelial dysfunction and progressively lead to more severe vascular alterations as the disease advances.

While our study represents an advancement in modeling PD-associated vascular pathology, there remain opportunities for further refinement and exploration. Notably, pericytes and microglial cells play essential roles in mediating inflammatory responses within perivascular regions, contributing to adverse vascular changes^40,112–115^. However, their absence in our current model suggests that the inflammatory mechanisms driving vascular alterations in PD may not be fully recapitulated. To address this, future efforts are necessary to advance our vascularized midbrain model by incorporating brain pericytes and microglial cells alongside endothelial and dopaminergic neuronal cells. This enhancement will provide a more pathophysiologically relevant model for investigating the complex cellular inflammatory interactions that contribute to PD-related vascular pathology. Additionally, although our model captures various aspects of PD-associated vascular pathology, it does not yet account for the angiogenic processes proposed to occur between BBB leakage and vascular regression^112,115–117^. Given the critical role of angiogenesis in vascular remodeling across various diseases^118–120^, this gap emphasizes the need for further studies to assess whether our engineering approach can accurately replicate the specific pathology of PD and fully emulate the multiple stages of PD-associated vascular alterations. Moving forward, investigating how α-synuclein fibrils regulate PD-associated angiogenesis could provide valuable insights into their broader impact on cerebrovascular remodeling. Furthermore, exploring whether vascular alterations arise directly from α-synuclein fibrils or indirectly through their effects on other molecular mediators represents an important direction for future studies using our vascularized midbrain model.

Despite these limitations, our study demonstrates the value of our vascularized midbrain model as a tool for emulating and investigating the abnormal vascular alterations implicated in the early stages and progression of PD which has been challenging with conventional in vitro approaches. Future improvements of this model should aim to incorporate a broader range of cellular interactions and pathophysiological responses to better mimic the complex dynamics of vascular alterations observed in PD.

### Conclusion

In conclusion, the study described here introduces a 3D vascularized midbrain model that represents a significant advancement in our ability to engineer the capillary interface of dopaminergic neurons in the SNpc. Upon exposure to PD-associated α-syn PFFs, this model successfully recapitulates key pathological features of PD, including α-synuclein aggregation, neuroinflammation, and progressive dopaminergic neuronal loss. Importantly, the model also enables the investigation of progressive PD-associated vascular impairments, revealing α-synuclein-induced endothelial dysfunction, barrier disruption, and vascular regression, which have previously been challenging to recapitulate using conventional in vitro technologies. These findings demonstrate that our engineering approach provides a promising PD model for probing specific disease mechanisms underlying the temporal dynamics of vascular pathology and its impact on neurodegeneration. Despite its strengths, we acknowledge that the absence of key supportive glial cell types such as pericytes, astrocytes, and microglial cells, which play integral roles in maintaining neuronal function and vascular integrity, may limit the translational relevance of our model. Future refinements should incorporate these cellular components and further mimic the structure and function of the native neurovascular unit to enhance physiological accuracy and pathophysiological relevance. Additionally, investigating the angiogenic responses associated with PD-related vascular alterations may provide further insights into disease progression.

Overall, our engineered model represents a substantial step forward in emulating key pathological features of PD, bridging significant gaps in current modeling approaches, and laying a solid technical foundation for advancing mechanistic studies. By enhancing our understanding of PD’s complexities, this platform promises to accelerate the development of therapeutic interventions specifically designed to address the dynamic interplay of the disease’s diverse facets.

## Methods

### Device Fabrication and Preparation

We used standard soft-lithography techniques to fabricate a poly(dimethylsiloxane) (PDMS) slab with recessed microchannels and reservoirs on both its top and bottom surfaces (**Fig. 1d**), which served as the top part of our device. For the bottom part, we also produced a plain PDMS slab following the same microfabrication scheme (**Fig. 1e**). As illustrated in **Fig. S1**, the cross-sectional dimensions of the neuron chamber, vascular channel, and side channels were 2 mm (width) × 700 μm (height), 1 mm (width) × 700 μm (height), and 2 mm (width) × 700 μm (height), respectively. The diameters of the opening in the neuron chamber and each were 2 mm and 5 mm, respectively. For fabrication of these components, PDMS (Sylgard 184, Dow Corning) base was mixed with a curing agent at a weight ratio of 10:1 (base:curing agent)and poured onto 3D-printed masters (Protolabs) (**Figs. d, e**). After degassing, PDMS was fully cured in an oven maintained at 55°C. The hardened PDMS was then removed from the molds, and inlet and outlet ports were created at both ends of the vascular channel and the two side channels in the microstructured slab to gain fluidic access. For device assembly, the microstructured slab was stamped onto a thin layer of uncured PDMS, which was prepared by spin-coating at 2500 rpm for 2 min, and then sealed against the plain PDMS slab that was used for the bottom part of the device. The assembled device was then baked at 55 °C to fully cure the PDMS adhesive.

Prior to cell culture, the assembled device was first sterilized by autoclaving with high-temperature pressurized steam (EZ Plus, Tuttnauer) for 1 hour. Subsequently, the neuron chamber, vascular channel, and side channels were filled with a 2 mg/mL polydopamine (PDA) solution (Sigma, H8502) prepared in 10 mM Tris-HCl (pH 8.5) to create a PDA coating that enhances extracellular matrix (ECM) hydrogel and cell adhesion to the PDMS surfaces within the device. After 2 hours of incubation at room temperature (RT), the PDA solution was gently aspirated, and the PDMSsurfaces were washed twice with sterile-filtered deionized (DI) water to remove any unbound PDA molecules. The washed device was kept sterile at RT until use.

### Cell Culture

To generate 3D vascular network in our device, we used primary human umbilical vein endothelial cells (HUVECs; C2519A, Lonza) and primary normal human lung fibroblasts (NHLFs; CC-2512, Lonza). HUVECs and NHLFs were cultured in 75 cm^2^ flasks following the manufacturer’s protocols. These cells were cultured and maintained using endothelial cell growth medium (EGM)-2 (CC-3162, Lonza) and fibroblast growth medium (FGM)-2 (CC-3132, Lonza), media respectively. HUVECs and NHLFs between passage 2 and 5 were used for device culture.

To establish neuronal cell culture within our device, the human neuroblastoma cell line (SH-SY5Y; ATCC, CRL-2266) was selected and expanded using 25 cm^2^ flasks according to the manufacturer’s protocols. To culture SH-SY5Y cells, Dulbecco’s Modified Eagle Medium (DMEM; 10-017-CV, Corning) was used as the basal growth medium, supplemented with 10% (v/v) heat-inactivated fetal bovine serum (hiFBS; SH30396.03, Cytiva), 1% (v/v) GlutaMAX-I (35050-06, Thermo Fisher) and 1% (v/v) penicillin-streptomycin (SV30010, Cytiva). SH-SY5Y cells at passages 3 to 5 were used for device culture, as recommended by the cell supplier.

### Vasculogenic Cell Culture

To form a vascularized tissue scaffold in our device, trypsinized HUVECs and NHLFs were resuspended in their growth media and mixed with fibrinogen (10 mg/mL; F8630, Sigma), thrombin (2 U/mL; T7513, Sigma), and aprotinin (1 U/mL; A1153, Sigma). The final density of HUVECs and NHLFs were 2.5 × 10⁶ cells/mL. The cell-containing hydrogel solution (20 μL) was then injected into the vascular channel, followed by incubation at 37°C with 5% CO₂ for 15 min to allow thermal gelation. Once the fibrin gel solidified, EGM-2 was introduced into the top medium reservoirs and the side channels to support vasculogenesis.

### Neuron Cell Culture

One day after the onset of vasculogenic cell culture, SH-SY5Y cells were trypsinized and resuspended in their growth medium. These cells were then introduced into the central neuron chamber of the device at a density of 2.5 × 10^5^ cells/mL. The initial culture was maintained in their growth medium to promote cell adherence and stabilization. One day after seeding, the SH-SY5Y cell growth medium was replaced with the first differentiation medium consisting of DMEM supplemented with 2.5% (v/v) hiFBS, 1% (v/v) GlutaMAX-I, 1% (v/v) penicillin-streptomycin, and 10 μM retinoic acid (50-165-6969, Fisher Scientific) to induce differentiation into dopaminergic neuronal cells. After 5 days, the first differentiation medium was switched to a second differentiation medium based on Neurobasal-A (10888022, Thermo Fisher) mixed with 50 ng/mL brain-derived neurotrophic factor (BDNF; P23560, FUJIFILM Irvine Scientific), 20mM potassium chloride (KCl; 7447-40-7, Avantor Science), 1% (v/v) B27 (17-504-044, Fisher Scientific), 1% (v/v) GlutaMAX-I, and 1% (v/v) penicillin-streptomycin. The second differentiation medium was also maintained for an additional five days. Both the first and second differentiation media were refreshed every other day to support and sustain the dopaminergic differentiation of SH-SY5Y cells.

### Production of PD-associated α-synuclein preformed fibrils

To generate α-synuclein preformed fibrils (α-syn PFFs), we replicated and applied the membranous lipid composition of neurons affected by PD, as this intraneuronal environment has been suggested to modulate the pathological properties of α-syn fibrils during disease progression. Specifically, the PD-associated α-syn PFFs were generated following the multistep procedure described below.

#### Protein Expression

Recombinant full-length α-synuclein was expressed in Escherichia coli (E. coli) BL21 (DE3) competent cells. These cells were grown in 1 L of Luria-Bertani (LB) medium supplemented with ampicillin (100 mg/mL) and incubated at 37 °C with 250 rpm until the optical density (OD600) reached 0.7-0.9. Protein expression was induced by adding 1 mM isopropyl-b-thiogalactopyranoside (IPTG). After 4 hours, the cells were centrifuged at 7,000 rpm for 20 min and stored at −80 °C.

#### Purification of α-Syn Peptides

The cell pellet obtained from 1 L of culture was resuspended in 25 mL of lysis buffer (10 mM Tris, pH 7.8, 1 mM EDTA, 1 mM PMSF, Sigma Fast protease inhibitor cocktail tablet), then lysozyme (Final concentration 0.2 mg/mL) was added and incubated in the ice bath for 20-30 min. Sonication was performed at pulse rate of 10-sec of on-time and 15-sec of off-time, with 40% amplitude power for total on-time of 3.5 min (Branson Digital Sonifier SFX 250, with Sonifier Sound Enclosure). Once the sonication is completed, the centrifuge tube is placed back into the ice bucket for 10-15 min before centrifuging at 4 °C with 16,000 rpm for 1 hour. The milky supernatant was collected and its pH was reduced to 3.5. The mixture was stirred at room temperature for 20 min and then centrifuged at 16,000 rpm for 30 min at 4 °C, resulting in a clear supernatant. The pH was then adjusted back to 7.5 and centrifuged at 16,000 rpm for 1 hour at 4 °C. The collected supernatant was passed through a PVDF, 30 mm syringe filter with 0.45 μm pore size (Avantar, VWR® Syringe Filters) and dialyzed against 1 L of low salt concentration anionic exchange buffer (10 mM Tris, pH 7.6, 25 mM NaCl, 1 mM EDTA) overnight at 4 °C in a dialysis membrane of MWCO 3.5 kDa (Spectrum Laboratories, Inc., Spectra/Por®3 Dialysis Membrane).

The dialyzed α-synuclein was filtered through a 0.25 mm PES syringe filter with a 0.22 μm pore size (Avantar, VWR® Syringe Filters) and loaded onto a double-stacked pre-packed HiTrapTM Q Sepharose High Performance anion exchange chromatography column (Cytiva Sweden AB). The column was washed with Buffer A (10 mM Tris, pH 7.6, 25 mM NaCl, 1 mM EDTA) and Buffer B (10 mM Tris, pH 7.6, 1 M NaCl, 1 mM EDTA). α-synuclein-containing fractions were pooled at conductivity of 30 mS/cm and concentrated to approximately 13 mg/mL using a 10 kDa Amicon ®Ultra centrifugal filter device (Millipore). The concentrated protein was loaded onto a pre-packed HiPrep 16/60 Sephacryl S-200 High Resolution preparative size exclusion chromatography column (Cytiva Sweden AB). Size exclusion chromatography was run in a buffer (10 mM Tris, pH 7.6, 100 mM NaCl) with a total 126 mL volume. α-synuclein protein eluted between 40 mL and 60 mL. All column chromatography was performed on an AKTATM pure system (GE Healthcare Bio-Sciences AB, Sweden). Protein concentration was determined using a NanoDrop ND-1000 UV/Vis Spectrophotometer (ThermoFisher Scientific). Purity was determined by gel electrophoresis. Purified α-synuclein is dialyzed against 1L of distilled water overnight at 4 °C and stored at −80 °C.

#### Preparation of PD-affected neuronal membranes

To mimic pathophysiologically relevant PD-affected neuronal membranes, 1,2-dipalmitoyl-sn-glycero-3-phosphocholine (DPPC), 1-palmitoyl-2-oleoyl-sn-glycero-3-phosphoethanolamine (POPE), cholesterol, and sphingomyelin (SM) were purchased from Avanti Polar Lipids. The PD-associated neuronal membrane composition consists of DPPC/POPE/cholesterol/SM at a molar ratio of 35:20:35:10. To prepare these membranes, phospholipid, cholesterol, and SM were co-dissolved in a chloroform/methanol mixture, then the solvent was evaporated under N_2_ gas. The remaining lipid film was then vacuum-desiccated overnight to remove residual solvent. The dried lipid films were resuspended to a final concentration of 12.5 mM in 12 mM Tris-HCl buffer (pH 7.6) and sonicated for 10-15 sec. The solution was then subjected to 10 freeze-thaw cycles using liquid nitrogen and a 50°C water bath, respectively, to generate homogeneous vesicles.

#### Fibril and α-syn PFFs Formation

Fibrils were generated by incubating 200 µM α-synuclein in 10 mM Tris-HCl buffer (pH 7.6) for four weeks at 37 °C without agitation in the presence of lipid vesicle solutions at a lipid-to-protein (L:P) ratio of 10. To prepare α-syn PFFs, the fibrils were broken using the sonifier (Branson Digital Sonifier SFX 250, with Sonifier Sound Enclosure) at a pulse rate of 1-sec of on-time and 4-sec of off-time, 10% amplitude power, and total on-time of 2 min. The average length of the sonicated α-syn PFFs (∼100 nm) was confirmed by negative-stained transmission electron microscopy and the prepared α-syn PFFs were stored at −80C.

### Transmission Electron Microscopy of α-syn PFFs

Negatively stained α-syn PFFs were prepared on 300-mesh lacey carbon-coated copper grids (Electron Microscopy Sciences, Ultrathin Carbon Films, LC-325-CU-CC). A 10 μL aliquot of the PFF solution was deposited onto the grids and allowed to absorb for 1-2 min. Subsequently, the excessive solution was blotted, followed by a wash with 10 µL of water and then blotted again and stained with 10 µL of UranyLess EM stain solution (Electron Microscopy Sciences) for 30 seconds. TEM images were acquired using a JEOL 2100F Field-emission transmission electron microscope at 120 kV, equipped with a Gatan Ultrascan CCD camera. The obtained images were processed using DigitalMicrograph (GMS3) software (Gatan Inc.).

### Treatment of PD-associated α-synuclein preformed fibrils

To investigate the pathological impacts of α-synuclein fibrils, our vascularized midbrain model was exposed to the PD-associated α-syn PFFs. For this work, the 200 µM stock solution of the PD-associated α-syn PFFs was first sonicated for 15 sec to ensure their uniform distribution and optimal working size (∼100 nm).

Subsequently, the stock solution was diluted in EGM-2 medium to a final concentration of 5 μg/mL for investigating its impact on the vascular component. Similarly, for treating the dopaminergic neuronal cells, the sonicated stock solution was diluted in the second dopaminergic differentiation medium to a final concentration of 1 μg/mL. After 11 days of cell culture within the device, the respective media containing the PD-associated α-syn PFFs were introduced into both the neuron chamber and side channels. The device was then incubated with the fibrils for 48 hours. Following the incubation, the media containing the PD-associated α-syn PFFs were replaced with fresh media in both the neuron chamber and side channels. The cell culture within the device was then maintained for an additional 3 days before analysis.

### ThioflavinT (ThT) assay for intraneuronal α-synuclein fibrilization analysis

To assess the extent of fibrilization induced by the treatment with the PD-associated α-syn PFFs within the dopaminergic neuronal cells, we performed the Thioflavin T (ThT) assay. A working solution of ThT was prepared in DPBS to a final concentration of 10 µM. This ThT solution was then added to both the neuronal chamber and the two side channels of the device which was subsequently incubated at RT for 10–15 min in the dark to prevent photobleaching. After incubation, the device was washed thoroughly twice with DPBS to remove any unbound ThT. Following this, the vessels and neuronal cells within the device were imaged using an inverted fluorescence microscope (Eclipse Ti2, Nikon). To generate quantitative data, the fluorescent intensity was averaged from 3 devices for each experimental condition.

### Measurement of Reactive Oxygen Species (ROS) production and Caspase-3/7 activation

Following the treatment with the PD-associated α-syn PFFs, the CellROX Green (5 μM in DPBS; C10444, ThermoFisher) and CellEvent Capase-3/7 Red (5 μM in DPBS; C10430, ThermoFisher) were used to measure oxidative stress and apoptosis of cells in our device, respectively. The dye solutions were injected into the device through the neuronal chamber and the two side channels and incubated at 37°C for 30 min. After three washing steps using DPBS, the vessels and neuronal cells were imaged using an inverted fluorescence microscope (Eclipse Ti2, Nikon). To quantitively assess the data, fluorescent intensity was averaged from 3 devices for each experimental condition.

### Immunostaining and Quantification

For immunostaining, cells within our device were fixed with 4% paraformaldehyde (AR1068, Boster) for 30 min at RT and washed twice using DPBS. After fixation, dopaminergic neuronal cells were permeabilized with 0.1% Triton X-100 (9002-93-1, VWR) for 15 min, while the permeabilization for vascular cells was extended to 30 min. To prevent non-specific binding, the cells were blocked with 1.5% bovine serum albumin (BSA; 22013, Biotium) for 30 min at RT. Subsequently, the cells were incubated overnight at 4°C with primary antibodies that selectively bind to the proteins of interest.

For imaging of the self-assembled vessels, we used a rabbit polyclonal anti-CD31 antibody (ab28364, 1:300-1:500, Abcam). To visualize the actin cytoskeleton, Phalloidin-CF 430 (Biotium) was applied at a dilution of 1:100. Additionally, neuronal differentiation within our device was demonstrated using primary antibodies of rabbit polyclonal anti-MAP2 (A16829, 1:200, Antibodies.com) for dendritic extensions, mouse monoclonal anti-GAP43 (A85392, 1:1000, Antibodies.com) for axonal projections, and mouse monoclonal anti-β III Tubulin (A86691, 1:500, Antibodies.com) for the neuronal cytoskeleton. To confirm PD-specific misfolding of endogenous α-synuclein, we used a rabbit monoclonal antibody against α-synuclein phosphorylated at Ser129 (A304933, 1:500, Antibodies.com). To assess the dopaminergic characteristics of neuronal cells, Tyrosine Hydroxylase (TH) expression was examined using a mouse monoclonal anti-TH antibody (A104316, 1:5000, Antibodies.com). For investigating inflammatory responses, we used rabbit polyclonal anti-NF-kB (51-0500, 1:500, ThermoFisher) and mouse monoclonal anti-ICAM-1 (A85684, 1:500, Antibodies.com) antibodies. For analyzing vascular endothelial integrity, we used a mouse monoclonal anti VE-cadherin antibody (14-1449-82, 1:100, ThermoFisher).

After incubation with primary antibodies, the cells were washed twice with DPBS and incubated for 1-2 hours at RT with fluorescently labeled secondary antibodies (A32732, ThermoFisher; A32723, ThermoFisher; A32731, ThermoFisher; A11032, ThermoFisher). We also used Hoechst (33342, ThermoFisher) for nuclear staining.

Finally, fluorescence images of the cells were captured using the Nikon Ti2-Eclipse microscope, and image processing was performed using NIS Elements software. To quantitively assess the data, fluorescent intensity was averaged from 3 devices for each experimental condition. For the quantitative assessment of VE-Cadherin expression, in particular, we measured the immunofluorescent intensity levels of VE-Cadherin in both healthy and unhealthy blood vessel areas for each sample. We then calculated the average intensity from these measurements to represent the expression level of VE-Cadherin in each sample.

### Statistical analysis

Statistical value of the obtained data was assessed by a two-tailed t-test. Data from three independent experiments were presented as mean ± SD.

## Acknowledgements

We thank S Choi for his input. This work was supported by the National Institutes of Health (NIH) (grant no. 1R21NS139178-01); Binghamton University (grant nos. TAE 1182867, ADLG258).

## Author contributions

A.A. and J.P. designed the research, performed the experiments, analyzed the data and wrote the manuscript. Y.B. and M.L. produced short form α-synuclein fibrils and wrote the manuscript.

