## Supplemental Figure 1 for "Microphysiological engineering of the capillary interface of substantia nigra dopaminergic neurons to study vascular alterations in Parkinson’s Disease"

**Supplementary Information (SI)**

**Supplementary Figure 1. Device dimensions. a,** The diameters of the opening in the neuron chamber and each media reservoir are 2 mm and 7 mm. **b,** The cross-sectional dimensions of the neuron chamber, vascular channel, and side channels are 2.0 mm (width) × 0.7 mm (height), 0.8 mm (width) × 0.7 mm (height), and 1.5 mm (width) × 0.7 mm (height), respectively.

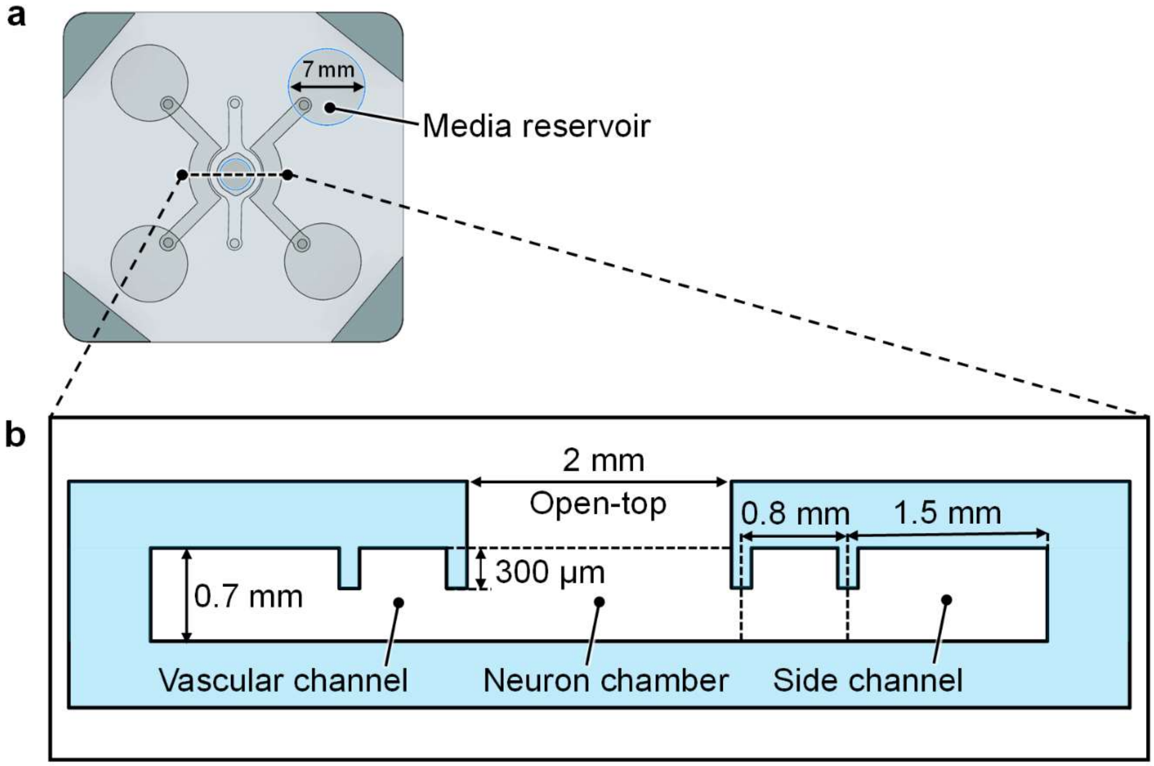
